# An extraglomerular relay circuit for multimodal integration within the Drosophila antennal lobe

**DOI:** 10.64898/2026.06.22.733763

**Authors:** Beate Bergkirchner, Rashmit Kaur, Alexandra Grimm, Sigrid Ilgerl, Wolfgang Kallina, Ameya Kasture, Dominik Javorski, Thomas Hummel

**Affiliations:** Dept. for Neuroscience and Developmental Biology, Division of Molecular Neurobiology, Faculty of Life Sciences, University of Vienna, Djerassiplatz 1, 1030 Wien, Austria

**Keywords:** neural circuit, olfactory system, commissural pioneer interneurons, single cell labeling, multimodal integration

## Abstract

The central integration of sensory information within and between brain hemispheres is critical for efficient behavioral responses, but at which level of information processing multimodal integration occurs is poorly understood. In olfactory systems an array of receptor-specific synaptic glomeruli with corresponding projection neurons (PNs) form separate sensory channels within each hemisphere, which run in parallel with other sensory modalities to converge at higher brain regions. We recently identified a small cluster of commissural pioneer neurons (cPINs) in the Drosophila olfactory system, which controls the formation of bilateral sensory circuits to support interhemispheric integration at the first synaptic layer. Here we show that cPINs also mediate the integration of sensory channels by relaying class-specific input within the antennal lobe. Functional studies showed that medial cPINs converge olfactory amine/ammonia input with a class of non-olfactory PNs to trigger attraction. During olfactory circuit formation, growing cPINs specify separate dendritic input/output domains, which merge into distinct glomeruli of different sensory modalities. Mutant analysis of the Wnt5 pathway revealed that cPINs display an initial PN-related growth pattern, which becomes redirected to organize lateral relay between sensory channels. These results identified a small cluster of olfactory interneurons as a central coordinator for fast convergence of sensory information, providing a mechanistic model of neural circuit evolution.

## Introduction

Animals use olfactory cues to navigate through their environment locating food and mates while avoiding potential threats like toxins and predators (Ache and Young, 2005; Wyatt, 2014). Whether a specific odor triggers an attractive or repulsive response behavior is determined by a complex interplay of odor identity in a given environmental context and the animals’ physiological state (Steck et al., 2012; Root et al., 2011). Fast integration of multimodal sensory information supports efficient behavioral response pattern and increases animal fitness (Stein and Stanford, 2008; Angelaki et al., 2009). However, multisensory integration for valence coding takes place in higher brain regions several synapses downstream of sensory receptor activation (Fiala, 2007; Wilson, 2013). In insects, conserved central brain domains like lateral horn and mushroom bodies encode and integrate different sensory modalities together with animal’s internal state (Marin et al., 2002; Wong et al., 2002; Sosulski et al., 2011; Aso et al., 2014). This circuit design seems specifically challenging for fast-flying insects like Drosophila in a competitive environment, where chemical cues must be integrate with vision, mechano- and thermosensation for proper path integration (Currier and Nagel, 2020). On the other hand, the fly’s main sensory systems are particularly well suited for early integration, as receptors for olfaction, thermo-, and hygrosensation are all located in the third antennal segment and project through a common antennal nerve to adjacent glomerular domains within the antennal lobe (Sayeed and Benzer, 1996; Chown and Nicolson, 2004; Marin et al., 2020).

Here, we characterize the developmental profile of a set of early extending bilateral olfactory neurons, previously identified as commissural pioneer interneurons (cPINs; Kaur et al., 2019), which bridge the spatial segregation of odor and humidity processing centers to enable early multimodal integration. During development, cPINs innervate the early AL prior to sensory axon arrival and extend contralaterally across the midline. Disruption of cPIN development leads to a shift from bilateral to unilateral innervation of all olfactory sensory neurons, as well as other commissural interneurons, indicating a central role in establishing main circuit organization. To further investigate subsequent steps of cPINs development we followed their integration into the mature Drosophila olfactory circuit. In addition to their role in early bilateral integration of olfactory inputs, we found that cPINs contribute to the connectivity between olfactory, thermal, and humidity pathways—three modalities essential for the fly’s survival and reproduction. Adult circuit analysis revealed that cPINs receive sensory input through a main glomerulus, while their output targets an aglomerular neuropil in the posterior antennal lobe. Developmental studies further showed that cPINs establish distinct spatial trajectories for their pre- and postsynaptic domains, ensuring appropriate innervation of both glomerular and aglomerular regions. Consistent with this organization, behavioral experiments demonstrated early integration of olfactory information with temperature and humidity cues.

## Results

### Dendritic fields of cPINs are organized in glomerular and extraglomerular domains

In the adult antennal lobe (AL) of Drosophila, a cluster of bilateral cPINs shows distinct, non-overlapping dendritic arborizations into a lateral, dorsal and medial AL domain (Figure 1A-C; Kaur et al., 2019). Single cell analysis combined with the recently described Drosophila olfactory connectome (Xu et al., 2020) revealed that this cPIN cluster consists of sparse LNs from the v2-cluster (v2LN5 and v2LN31, Figure 1E) and a multiglomerular PN derived from the l2 lineage (l2PN^VP4+VL1^, Figure 1E). The dendrites of each cPIN class are associated with a specific sensory glomerulus: The bilateral LNs-type cPINs cover olfactory glomeruli, VM6 for cPIN^LN5^ and DP1m for cPIN^LN31^, while the bilateral l2PN contact posterior VP glomeruli of thermo/hygro modalities (Figure 1C, Suppl. Figures).

**Figure 1.**
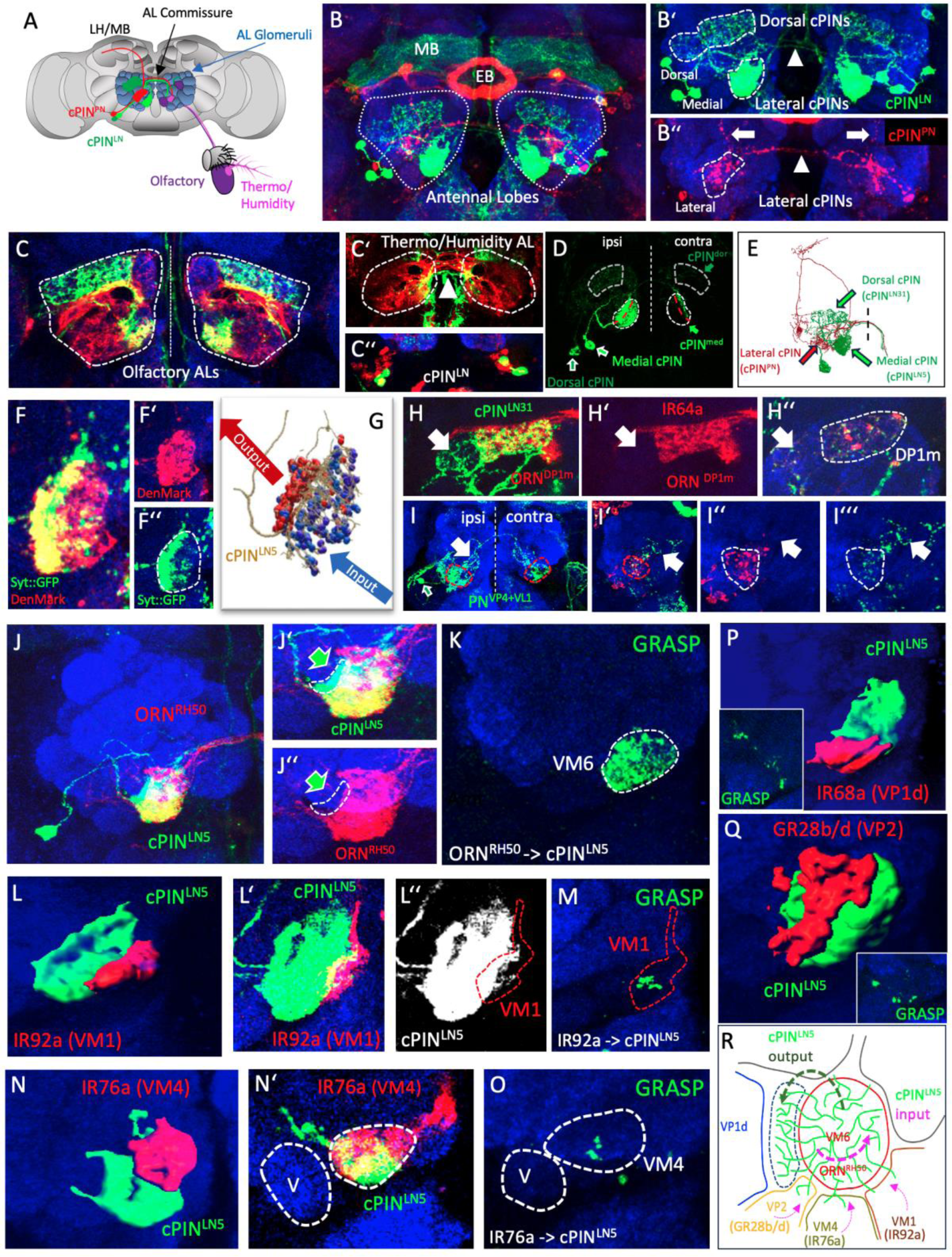
Organization of bilateral cPINs in the posterior antennal lobe. (A-A’’) Organization of antennal sensory systems in adult Drosophila. The 3^rd^ antennal segment contains mainly olfactory sensilla with separated sensory domains for temperature (arista) and humidity (sacculus). Antennal sensory axons segregate into distinct glomeruli in the anterior (olfaction) and posterior (temperature/humidity) antennal lobe (AL). Relay neurons of bilateral cPIN classes occupy distinct, non-overlapping AL areas and can be divided into local cPINs (cPIN^LN5^ in green) and relay cPINs (cPIN^PN^ in red). (B-B’’) Localization of cPIN dendrites in the posterior AL adjacent to the Ellipsoid Body (EB) of the central complex and the mushroom body (MB) lobes. Cell body position and dendritic pattern of medial and dorsal cPIN^LN5^ (B’) and lateral cPIN^PN^ (B’’, with dorsally-projecting axons, arrows) and commissural projections of both cPIN classes (arrowhead). (C-C’’) Multiple types of bilateral interneurons (anterior-ventral cell cluster in C’’) cover the posterior AL in the transition from defined olfactory glomeruli (C) to more variable glomeruli of thermo- and hygro modalities (C’, arrow indicates AL commissure). (D) Single cell analysis of medial and lateral cPINs illustrating the asymmetry in their ipsi-/contralateral dendritic arborization. (E) Matching cPIN class identity with cell types of the Drosophila hemibrain connectome. While both the medial and dorsal cPIN classes are subtypes of bilateral sparse local interneurons from the v2 lineage (v2LN05 and v2LN31 respectively), the lateral cPIN class matches the morphological organization of bilateral l2 PNs (l2PN^VP4+VL1^). (F-K) Domain organization of cPIN dendrites visualized with presynaptic Syt::GFP (green) and postsynaptic DenMark:RFP (red) markers, which correspond to the input/output regions from the hemibrain data set (G). Dorsal cPIN^LN31^ (H-H’’) and lateral cPIN^PN^ (I-I’’) also display restricted input regions (dashed lines) and more distributed presynaptic output processes (arrows). (J-Q) Colabeling of medial cPIN^LN5^ (green) with different sensory neuron classes (red). (J) The dendritic field of medial cPIN^LN5#^ show a strong colocalization with ORNs expressing the ammonia-receptor RH50 (red) except for a specific lateral region (dashed lines indicated by green arrows in J’ and J’’). (K) Transsynaptic GFP reconstitution (GRASP) supports the synaptic input of ORN^RH50^ onto medial cPIN^LN5^ within the VM6 glomerulus. (L-Q) Other sensory classes which target distinct AL glomeruli surrounding the VM6 glomerulus also show minor spatial overlap with medial cPIN^LN5^ dendrites including IR classes in VM1 (IR92a), VM4 (IR76a), VP1d (IR68a) and the GR68b/d class in the posterior VP2 glomerulus. Corresponding GRASP signal could be detected for all overlapping sensory classes indicating direct synaptic input onto medial cPIN dendrites. (R) Schematics illustrating the organization of medial cPIN^LN5^ dendrites (green fibers) with one major (RH50) and multiple minor olfactory input region and a restricted non-sensory output region.

Interhemispheric innervation of different cPINs is highly asymmetric in the bilateral coverage of the homotopic glomeruli, with a large dendritic field on the ipsilateral side and a sparse innervation in the contralateral hemisphere (Figure 1D). Although ipsilateral dendritic fields overlap with specific glomerular units, co-labeling of individual cPINs with corresponding olfactory sensory classes showed only a partial spatial match, in which cPIN dendrites extend beyond the glomerular boundaries defined by ORN axons and uniglomerular PN dendrites (Figure 1H, J). These extraglomerular dendritic regions can be rather restricted for medial cPIN^LN5^ (Figure 1J) or occupy a more extended region in the posterior AL seen for dorsal and lateral cPINs (Figure 1H).

The morphological class of sparse LNs often displays intrinsic polarity of their dendritic domains in separated input and output zones (Chou et al., 2010; Xu et al., 2020). Expression of the somato-dendritic marker DenMark::mCherry showed an even distribution throughout their dendritic fields, whereas the presynaptic markers (Syt::GFP/Brp::GFP) are strongly enriched in the extraglomerular region (Figure 1F). This spatial segregation of cPIN dendritic field is distinct from other central AL neurons in which presynaptic differentiation can be observed throughout the dendritic field (LN and PNs -> VM6; Nicolai et al., 2010; Mosca and Luo, 2014). Segregated input/output domain organization of cPINs is supported by directional trans-synaptic GFP reconstitution (GRASP, Figure 1K). ORN^RH50^ provides synaptic input to cPIN^LN5^ dendrites only in the glomerular domain and no sensory connections could be detected in the extraglomerular field (Figure 1K). Colabeling of medial cPINs with neighboring ORN classes showed that in addition to the main olfactory input by the VM6-specific ORN^RH50^ class, cPIN dendrites also contact sensory axons of adjacent olfactory glomeruli like VM1 (ORN^IR92a^), VM4 (ORN^IR76a^) (Figure 1L-O). Both ORN classes are tuned to amine-class odorants (Hussain et al., 2016). Furthermore, non-olfactory glomeruli along the posterior surface including VP2 (GR28b/d) and VP1d (IR68a) show spatial overlap with cPIN^LN5^ processes (Figure 1P, Q). Positive GRASP signals at the side of physical overlap for all sensory classes confirmed direct synaptic connections with medial cPINs, although to a much lesser extent compared to VM6 (Figure 1K, M, O; insets in P, Q). Taken together, polarized medial cPINs integrate sensory activity of different amine-specific ORN classes as well as minor input from sensory neurons for humidity and temperature (Figure 1R). With the presynaptic sites of cPIN dendrites located in the posterior AL, this cell type defines an interesting candidate for mediating directional communication between sensory channels.

### cPIN^LN5^ relays olfactory information to a non-olfactory output channel

To further define the extraglomerular circuit organization built by cPIN dendrites we characterized the spatial connectivity of medial cPINs with surrounding PN classes. Similar to the colabeling of cPIN^LN5^ with ORN^VM6^, a strong spatial overlap could be observed with the corresponding projection neurons (PNs) of VM6, defining the glomerular boundaries (Figure 2A-D). Also, for PN^VM6^ the lateral domain of cPIN^LN5^ dendrites which are devoid of sensory axons, are lacking any direct contact to PN^VM6^ dendrites (Figure 2E). However, this extraglomerular dendritic domain of cPIN^LN5^ colocalizes with PNs from the adjacent VP1d glomerulus (Figure 2F). GRASP analysis showed synaptic input from cPIN^LN5^ onto PN^VP1d^ but only minor intraglomerular connections onto PN^VM6^ (Figure 2K-M). While IR40a-positive sensory neurons of VP1d are not in contact with medial cPINs (Suppl. Figures), dendrites of PN^VP1d^ extend into the extraglomerular region to receive synaptic input from cPIN^LN5^. This spatial pattern defines the extraglomerular region of cPIN dendrites as a site of directional lateral relay within the AL (Figure 2S). Connectome data indicate two main PN classes for the VP1d glomerulus, which receive IR40a-positive sensory input, il2PN^VP1d^ and l2PN^VP1d+VP4^ and cover different glomerular domains (Figure 2S). While l2PN^VP1d+VP4^ colocalizes with IR40a neurons and even extending into the dorsal VA4 glomerulus, il2PN^VP1d^ dendrites remain restricted to the ventral region of VP1d but cross the glomerular border and overlap with the adjacent extraglomerular cPIN^LN5^ domain (Figure 2S). Interestingly, in addition to medial cPIN^LN5^, similar morphological classes of sparse LNs, vLN27,28/29, receive sensory input within VM6 to relay to il2PN^VP1d^ dendrites within the extraglomerular domain, either directly or via cPIN (Figure 2S). This circuit architecture indicates the early convergence of the olfactory ammonia channel with the sensory pathway for temperature perception via a distinct relay LN-PN pathway (Figure 2S).

**Figure 2.**
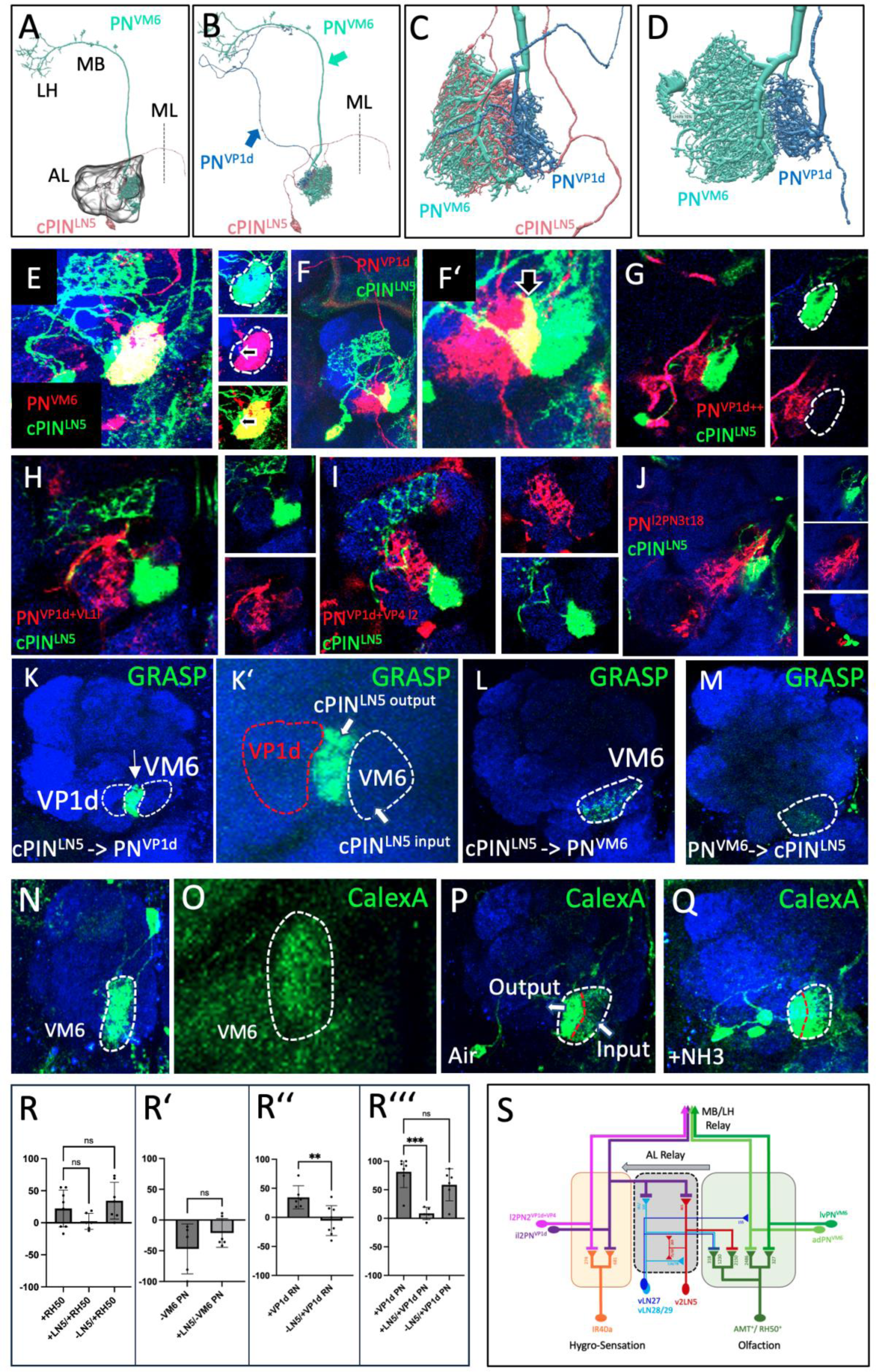
Medial cPIN^LN^ relay olfactory information to a non-olfactory output channel. (A-D) Dendritic domains at the transition from olfactory to non-olfactory AL regions, as represented in hemibrain. Medial cPIN^LN5^ dendrites are spatially overlapping with unilateral PNs of the two adjacent glomeruli VM6 and VP1d, with different relay pathways towards MB and LH (arrows in B). While dendrites of the two PN classes cover adjacent AL domains but do not spatially overlap (D), dendrites of the medial cPIN^LN5^ class overlap with the dendritic fields of both PN classes to different degrees (C): completely with PN^VM6^ and partially with PN^VP1d^. (E) Colabeling of the main uniglomerular PN class of the VM6 glomerulus (red) with cPIN^LN5^ (green) shows a similar overlap as ORN^VM6^, further supporting a defined extraglomerular domain of cPIN^LN5^ dendrites (arrow). (F, F’) Colabeling of an uniglomerular PN class for the VP1d glomerulus (red) with cPIN^LN5^ (green) shows a corresponding spatial overlap within the extraglomerular domain of cPIN^LN5^ dendrites (arrow in F’). (G) A second PN class of the VP1d glomerulus with a dendritic field restricted to the glomerular boundaries are spatially separated from cPIN^LN5^ dendrites (dotted line). (H-J) Multiglomerular VP1d classes adjacent to the extraglomerular cPIN^LN5^ dendrite domain show no spatial colocalization, indicating a specific relay from VM6 to VP1d via cPIN^LN5^. (K-M) GRASP-mediated GFP expression support strong extraglomerular dendro-dendritic cPIN^LN5^-> VP1d connections (K) and a small amount of intraglomerular cPIN^LN5^-> VM6 synapses (L). (M) Within the VM6 glomerulus, a weak GRASP signal could be detected for PN^VM6^ onto cPIN^LN5^ dendrites, indicating a directional segregation of olfactory input into an intra- and extra-AL relay. (N-Q) Ca^2+-^-induced GFP expression revealed distinct soma-dendritic Ca2^+^ levels for PN^VM6^ onto cPIN^LN5^: a low GFP increase in PN^VM6^ following ORN activation (N, O), compared to the high-default GFP expression in cPIN^LN5^ which can be further increased by sensory stimulation (P, Q). (R) Optogenetic manipulation of cPIN^LN5^ in the context of VM6/VP1d. cPIN^LN5^ reduces the ORN^VM6^ induced attraction (R) but has little effect on the ORN^PN^ mediated repulsion (R’). For the VP1d pathway, cPIN^LN5^ silencing blocks the sensory receptor RN^VP1d^ induced attraction (R’’) while the activity of cPIN^LN5^ blocks the attraction by PN^VP1d^ (R’’’). (S) Schematics of cPIN^LN5^ circuit organization within two synaptic glomeruli of different sensory modality. Designated sensory neurons for olfaction (RH50) and hygrosensation (IR40a) synapse onto distinct PN classes (green and purple, respectively) within adjacent glomeruli (VM6, VP1d) in the posterior AL. In addition to the PN-mediated relay of glomerulus-specific activity towards the Lateral Horn (LH) and Mushroom Body (MB), the medial cPIN^LN5^ class (v2LN5) relays olfactory input from VM6 onto il2PN^VP1d^ PNs within an extraglomerular domain (grey box), thereby merging different sensory modalities. Based on HB connectomics data, two additional classes of sparse local interneurons (vLN27, vLN28/29) support v2LN5 lateral relay. The number indicated weighted synaptic input (HB).

To gain insights into the role of cPINs in olfactory processing we analyzed cPIN^LN5^ activity upon sensory stimulation. Olfactory activation of the ammonium-sensitive RH50 ORNs leads to a strong increase in Ca^2+^ levels in cPIN^LN5^ by an elevated CaLexA signal (Figure 2N-Q). Here, the highest level of the Ca^2+^ sensor can be detected in the presynaptic extraglomerular dendrite domain (Figure 2N, O). In contrast, only a subtle CaLexA signal is induced within the glomerular relay neuron PN^VM6^ (Suppl. Figures), indicating different modes of activation patterns between PN^VM6^ and the cPIN^LN5^ pathways. No significant level of CaLexA signal can be detected for IR40a-positive sensory neurons innervating the VP1d glomerulus following ammonium application (Suppl. Figures).

To test the role of cPIN^LN5^ in the interaction of the VM6 and VP1d glomeruli, we determined adult fly behavioral responses following optogenetic activation of distinct neuron types (Figure 2R). We first examined the fly’s innate response to selective activation of cPIN^LN5^. This manipulation produced a completely neutral behavior, with no preference for either the illuminated or the dark side of the experimental arena (Suppl. Figures). Similar to the sensory stimulation of RH50-positive ORNs by ammonia (Vulpe et al., 2021), the expression of the red-activatable Channel-Rhodopsin ReaChR (Inagaki et al., 2014) in ORN^RH50^ resulted in a moderate attraction of flies towards the corresponding light source (Figure 2R). Interestingly, co-activation of RH50-positive ORNs and cPIN^LN5^ interneurons show a trend of suppressing the sensory-induced attraction (Figure 2R), though this effect did not reach statistical significance, suggesting that the independent activation of the two main relay channels of ammonia-induced attraction is not sufficient to trigger the same response but rather an antagonistic function of LN5 of ammonia-induced attraction.

Optogenetic activation of the main uniglomerular projection neuron PN^VM6^ is not sufficient to elicit attraction; instead, it induces a mild aversive response (Suppl. Figures). This aversive behavior becomes even more pronounced when PN^VM6^ is optogenetically silenced. These results suggest that PN^VM6^ does not directly mediate ORN_RH50-driven attraction but rather functions to suppress the fly’s innate aversive response to ammonia (Silbering et al., 2011; Min et al., 2013). Activation of VP1d receptor neurons for hygro-/thermosensation (Marin et al., 2020) elicits a moderate attraction (Fig 2R’’). While co-activation of VP1d receptor neurons with cPIN^LN5^ was not possible due to lethality, the inactivation of cPIN^LN5^ suppresses VP1d -mediated attraction (Figure 2R’’) (Marin et al., 2020). Similarly, the behavioral attraction induced by PN^VP1d^ activation is fully suppressed by the coactivation of cPIN^LN5^ (Fig 2R’’’). As no effect on VP1d -mediated attraction could be observed following the silencing of cPIN^LN5^, these interneurons seem to act as inhibitors of PN^VP1d^ -induced behavior (Figure 2R’’’)

### cPIN subtypes share a common bilateral growth program

To gain further insights into the intra- and extra-glomerular organization of cPINs within olfactory circuits we analyzed their developmental profile in the context of ORN/PN assembly. In late 3rd instar larvae, about 20hrs before ORNs arrive at the newly forming adult AL, cPINs start to grow into the neuropil precursor in a class-specific temporal sequence (Figure 3A). The lateral cPIN^PN^ class innervates the ventral AL first and extends dorsally beyond the AL neuropil along the main PN fiber tract (Figure 3A, B). At the ventral border of the horizontal MB lobes, lateral cPINs pause their dorsal growth and extend a single commissural process across the midline (Figure 3B). cPIN commissures stall again at the dorsal region of the contralateral AL neuropil to contact the ipsilateral branch of the corresponding cPIN^PN^ counterpart (Figure 3C, D).

**Figure 3.**
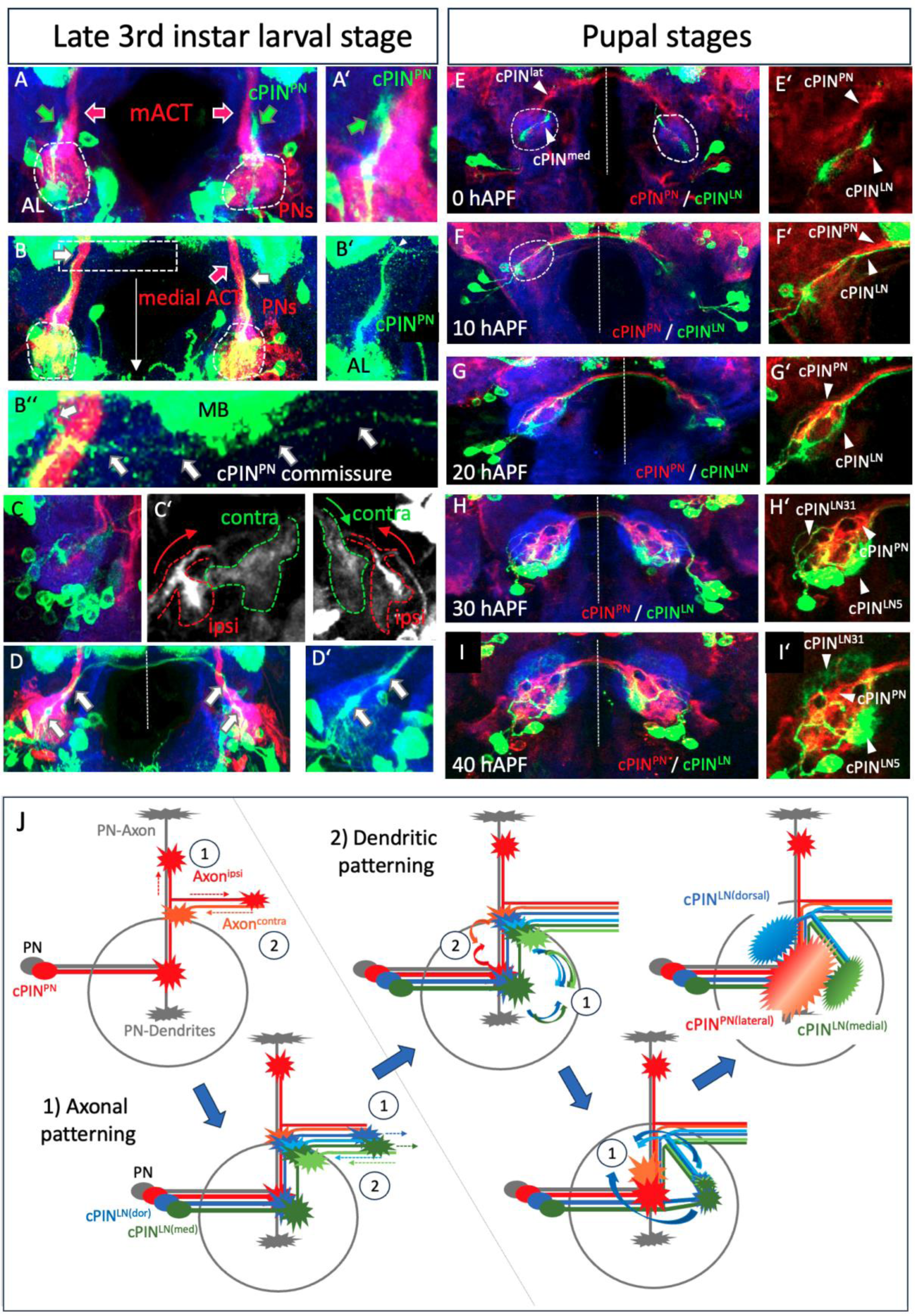
Developmental sequence of cPIN axonal and dendritic patterning. (A) At late 3rd instar larval stage, lateral PN-type cPINs (green arrows) extend along the existing medial axon tract of larval PNs (red arrows) to exit the AL (dotted line). (B-B’’) At the level of the horizontal MB lobes, cPIN^PN^ axons branch (white arrows) to send a thin commissural extension across the midline (B’’). (C-D) The ipsi- and contralateral cPIN axon branches (C’) converge in a dorsal domain, separated from the ventral dendritic field (white arrows). (E, F) In the first 10 hrs of pupal development, medial and dorsal LN-type cPINs are following the pioneer cPIN^PN^ projection within the AL and establish similar ventral and dorsal arborizations (E’). In contrast to the cPIN^PN^ class, cPIN^LN^ axons do not extend further along the PN tract but grow along the cPIN^PN^ commissural branch across the midline (F, F’). (G-H) Within the next 20 hrs, Ipsi- and contralateral branches of LN- and PN-cPINs converge into distinct regions (medial versus lateral, respectively) and increase in size to build adjacent, non-overlapping domains in the AL (H). (I, I’) Starting around 40hrs APF, the dorsal cPIN^LN31^ class extends from the medial domain around the cPIN^PN^ field dorsally to elaborate an additional cPIN domain, thereby establishing the cPIN spatial ground pattern of the adult AL. (J) Schematics illustrating the main developmental steps of cPIN growth and patterning with the ipsi- and contralateral growth (1 and 2, respectively) of cPIN^PN^ (red) followed by the joined extension of the two cPIN^LN^ classes (blue and green). Subsequent class-specific convergence of cPIN processes into distinct AL domains (1 and 2 for LN- and PN-types) before the cPIN^LN31^ type establishes a 3^rd^ domain in the dorsal AL.

About 5hrs later (0hrs APF), medial and dorsal cPIN^LN^ classes start growing along the cPIN^PN^ tract into the ipsilateral AL (Figure 3E). Following a short pause (about 5-10 hrs), individual cPIN^LN^ processes fasciculate with the established cPIN^PN^ commissure to project contralaterally (Figure 3F). In the next 20hrs of AL development, the ipsi- and contralateral cPIN branches segregate in a class-specific manner: First, medial and dorsal cPIN^LN^ dendrites converge together into the medial-ventral AL whereas lateral cPIN^PN^ occupy the central AL domain (Figure 3G, H). Secondly, cPIN^LN^ send a neuronal process from the ventral domain towards the dorsal AL (Figure 3H). By the time of AL glomerulus formation (30-40h APF), cPIN^LN^ dendrites innervate their class-specific glomerular domains in a non-overlapping fashion (Figure 3I). These results show that the different cPINs display a sequential but similar initial growth pattern in close association with PN dendrites to subsequently diverge in a class-specific manner and segregate into distinct glomerular domains.

### Patterning of cPIN axon branches in a defined spatial and temporal order

To characterize the unique polarized organization of cPIN dendrites we analyzed cPIN^LN5^ in single cell mosaics (Figure 4). Following initial growth into the ipsilateral AL, individual cPIN^LN5^ form two distinct branching domains, a proximal one at the ventral AL and a distal one at the dorsal AL edge (Figure 4A, B). These two branch domains differ in the distribution of the cell recognition molecule Dscam, which is highly enriched in the distal cPIN^LN5^ process (Figure 4C). Upon contralateral extension around 10hrs APF (Figure 4C), a filopodia-rich domain remains at the dorsal AL resulting in three growth-active regions of each cPIN^LN5^ (ipsi-ventral, ipsi-dorsal, contra-dorsal). While all cPIN^LN5^ commissural processes are stalling at the entry side of the contralateral AL, the two ipsilateral branch domains of cPINs start to converge into a common neuropil (Figure 4D, E). In the case of cPIN^PN^, this is a defined region in the central AL precursor (data not shown). For cPIN^LN5^, convergent growth is initiated by the medial extension of ipsi-ventral filopodia and followed a few hours later by the ventral growth of the ipsi-dorsal branches (Figure 4D). The contralateral axon is closely following the ventral path of the ipsi-dorsal branches and merges together with the two ipsi-lateral branches in a large medial protoglomerulus (Figure 4E).

**Figure 4.**
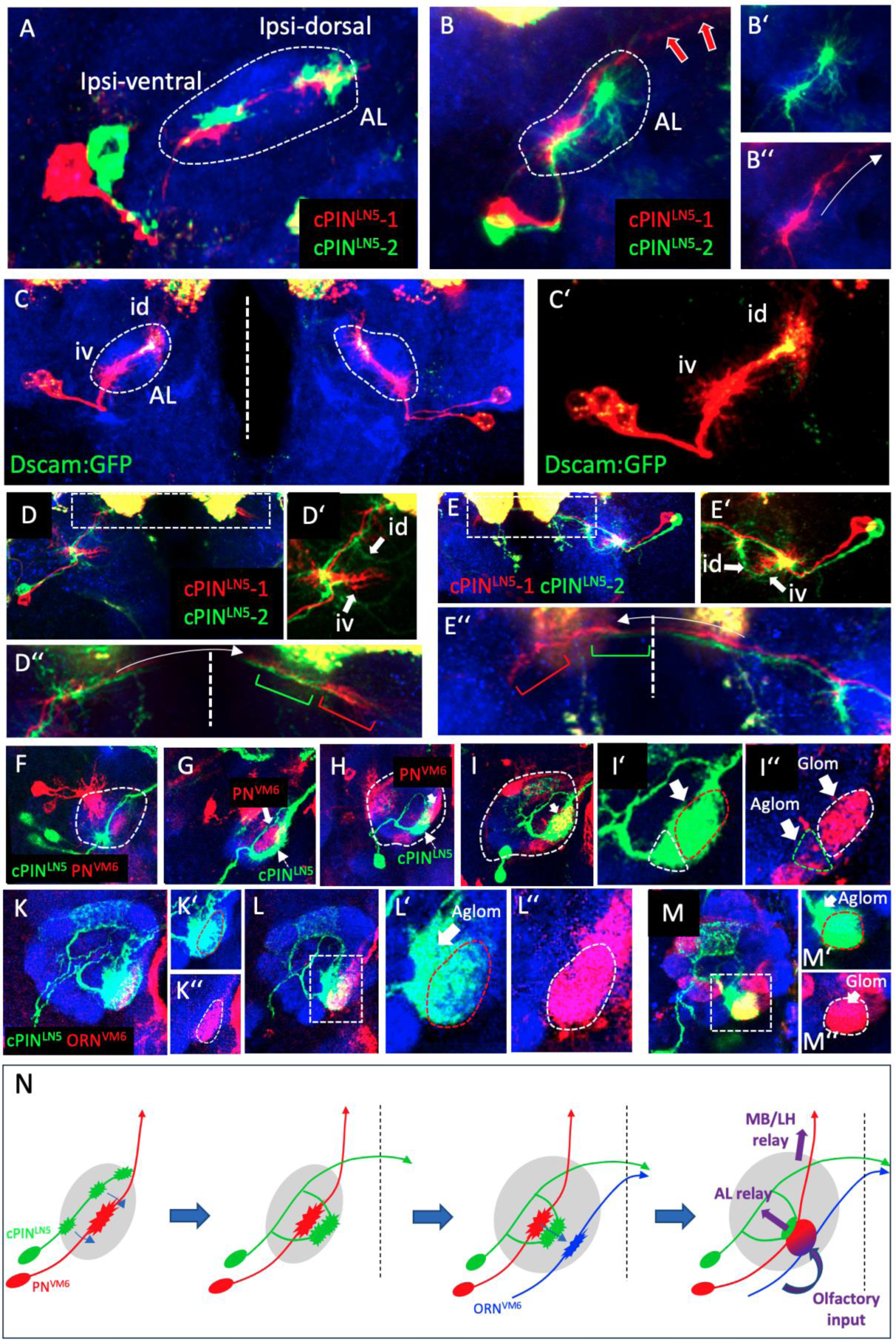
cPIN axon-dendrite dynamics in olfactory circuit assembly. (A, B) Upon initial dorsal extension of medial/dorsal cPIN^LN5^ the formation of two distinct ipsilateral branch sites (ventral and dorsal branch, iv/id) can be observed. While the ventral branch side remains stationary (B’), the dorsal branch side sends a contralateral branch extension across the dorsal midline but leaving a cellular mark at the position of the initial dorsal branch side (B’’). (C, C’) Differential branch dynamics illustrated with the enrichment of a transgenic Dscam:GFP at the dorsal branch side before contralateral extension. (D, E) Upon arrival of contralateral branches (D’’), ventral and dorsal branches send processes towards each other (arrows in D’) and meet and the ventro-medial AL surface (arrows in E’). (F-I) Sequential interaction of cPIN^LN5^ processes with PN dendrites. During ventral and dorsal branch convergence at the medial AL, cPIN^LN5^ processes grow around a cluster of PN dendrites (F, G) before contacting their future target glomerulus VM6 (H). In the following stage of glomerulus formation, the majority of cPIN^LN5^ processes integrate into the PN glomerular domain (I) except a ventral extraglomerular part (I‘ and II’’). (K-M) With the formation of protoglomeruli by ORN axons, (red in K’’), cPIN^LN5^ processes grow into the ORN/PN glomerulus (dotted lines) while a more condensed cPIN^LN5^ domain remain outside of the glomerular boundaries (arrow in M’). (N) Schematics illustrating the growth and branch patterning of cPIN^LN5^ before (1, 2) and after (3, 4) ORN axon ingrowth. cPIN^LN5^ processes first integrate into PN dendrites along the ventro-dorsal axis followed by medial-lateral extension for glomerular ORN axon integration.

Ipsilateral cPIN^LN5^ branch extension and convergence occur in synchrony with PN dendrite patterning and ORN axon innervation (Figure 4F-M). For ipsilateral extension, dorsal and ventral cPIN^LN5^ branches grow along the border of the early PN^VM6^ dendritic field, which becomes fully encircled upon ventral branch convergence (Figure 4F, G). In the next 10 hours, the dendritic fields of cPIN^LN5^ and PNs remain segregated but are swapping their relative position along the medial-lateral AL axis (Figure 4H). During ORN axon innervation, cPIN^LN5^ dendrites extend medially and merge with the PN dendritic field. In the following 10 hours of pupal development, the dendritic domains expand in a cPIN class-specific pattern (Figure 4I-M).

### Early segregation of pre-and postsynaptic domains within growing cPINs

To better understand the formation of distinct input/output domains of cPIN^LN5^ dendrites we followed the distribution of transgenic marker proteins Syt::GFP and DenMark::mCherry which in Drosophila segregate into pre- and postsynaptic compartments respectively (Nicolai et al., 2010). From the initial ipsilateral cPIN^LN5^ growth on, DenMark::mCherry enriches in the ventral branch domain whereas presynaptic Syt::GFP localizes to the dorsal domain, indicating an early axo-dendritic polarity (Figure 5A, B). During commissural branch extension, Syt::GFP is highly enriched in both the ipsi- and contralateral dorsal branches whereas DenMark::mCherry remains restricted to the ventral branch domain, which is now also accumulating Syt::GFP (Figure 5C). In the next 10 hours, both branch compartments will merge into a common elongated dendritic field in which presynaptic and postsynaptic domains are mainly overlapping (Figure 5D-H). During subsequent glomerulus formation/maturation, the ventral DenMark::mCherry positive dendrite domain extends into the newly-forming VM6 glomerulus without any signs of Syt::GFP (Figure 5I-K). These data indicate 2 phases of cPIN development in which early-specified extraglomerular dendritic domains are first organized within distinct AL regions which is followed by directional growth of their postsynaptic domains into distinct glomeruli (Figure 5L).

**Figure 5.**
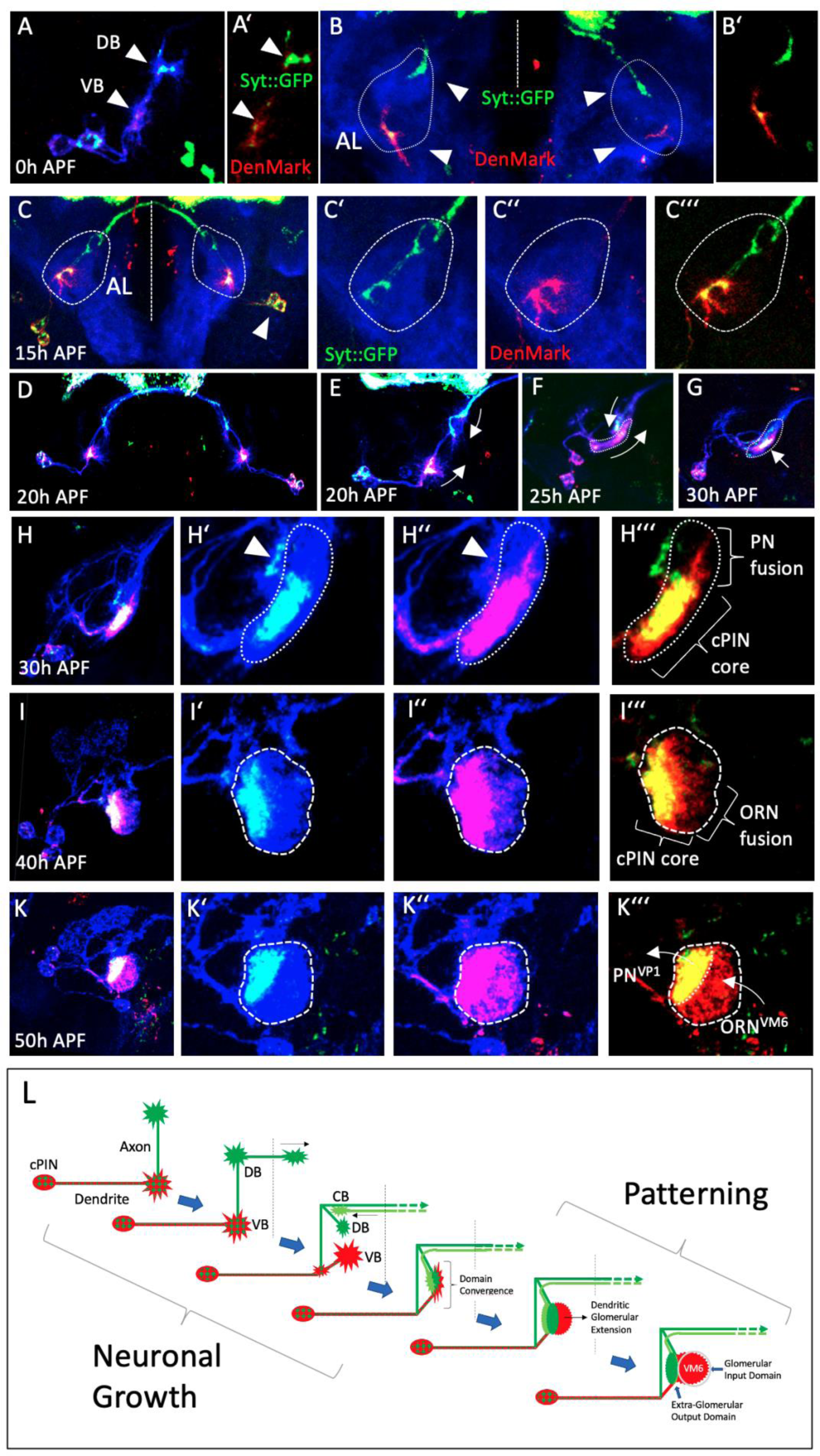
Early specification of cPIN input/output domains. (A-B) cPIN^LN5^ -specific expression of transgenic markers for presynaptic sites (Syt::GFP in green) and dendritic compartments (DenMark:RFP in red) reveals an early branch polarity in which the ventral branch (VB) constitutes dendrite identity whereas the dorsal branches (DB) accumulate presynaptic sites. (C-C’’’) At the beginning of cPIN^LN5^ ipsilateral branch convergence (arrowhead indicate cell bodies), an increase of presynaptic Syt::GFP can be observed at the ventral branch site. (D-G) Following cPIN branch convergence at the ventro-medial AL (arrows in E), a colocalization of the axonal presynaptic sides with the dendritic domain in a restricted “core domain” can be observed (arrow in G). (H-K) During glomerulus assembly, axonal and dendritic processes of the “core domain” show a different growth pattern. While dendritic processes of cPINs follow the sequential integration, first with dorsal PN dendrites (H-H’’’) and afterwards with lateral ORN axons (I-I’’’), the axonal domain remains stationary, thereby ending up in an extraglomerular region (K-K’’’). (L) Schematics illustrating the patterning of cPIN dendrites into distinct pre-/postsynaptic regions (VB ventral branch, DB dorsal branch, CB contralateral branch).

### Wnt signaling suppresses a default PN differentiation program to support bilateral cPIN organization

We previously identified the cell adhesion molecule Neuroglian (Nrg) in the regulation of cPIN interactions with ORN axons to support bilateral sensory circuit formation (Kaur et al., 2019). For further characterization of early cPIN patterning we analyzed additional mutations which result in *Nrg*-like phenotypes and identified *Wnt5*. In contrast to *Nrg* mutants, in which ORNs switch from a bilateral to a precise unilateral innervation pattern (Kaur et al., 2019), loss of *Wnt5* results in frequent ectopic projection of ORN axons in a dorsal direction (Wu et al., 2014; Hing et al., 2020). Overall pattern of cPIN projections in the adult AL is maintained in *Wnt5* mutants with a reduction in the dendritic fields and a loss of the commissural tract (Figure 6A, B). Developmental analysis revealed a corresponding defect in early cPIN^LN^ projection following the loss of *Wnt5* (Figure 6C-F). cPIN^LN^ in *Wnt5* mutants start to project into the ipsilateral AL like in wild type but show reduced ventral and dorsal branch domains (Figure 6C, D). In contrast to the horizontal extension of the dorsal branch to the contralateral AL, cPINs in *Wnt5* mutants fail to pause at the dorsal AL and follow the PN axon fascicle towards MB/LH regions (Figure 6E, F). Despite a redirection of the axonal branches, compartment-specific protein distribution in *Wnt5* mutants showed no change in cPIN^LN^ neuronal polarity with a proximal accumulation of DenMark::mCherry together with Syt::GFP and Syt::GFP-only distal process which extends dorsally instead crossing the midline (Figure 6G-I).

**Figure 6.**
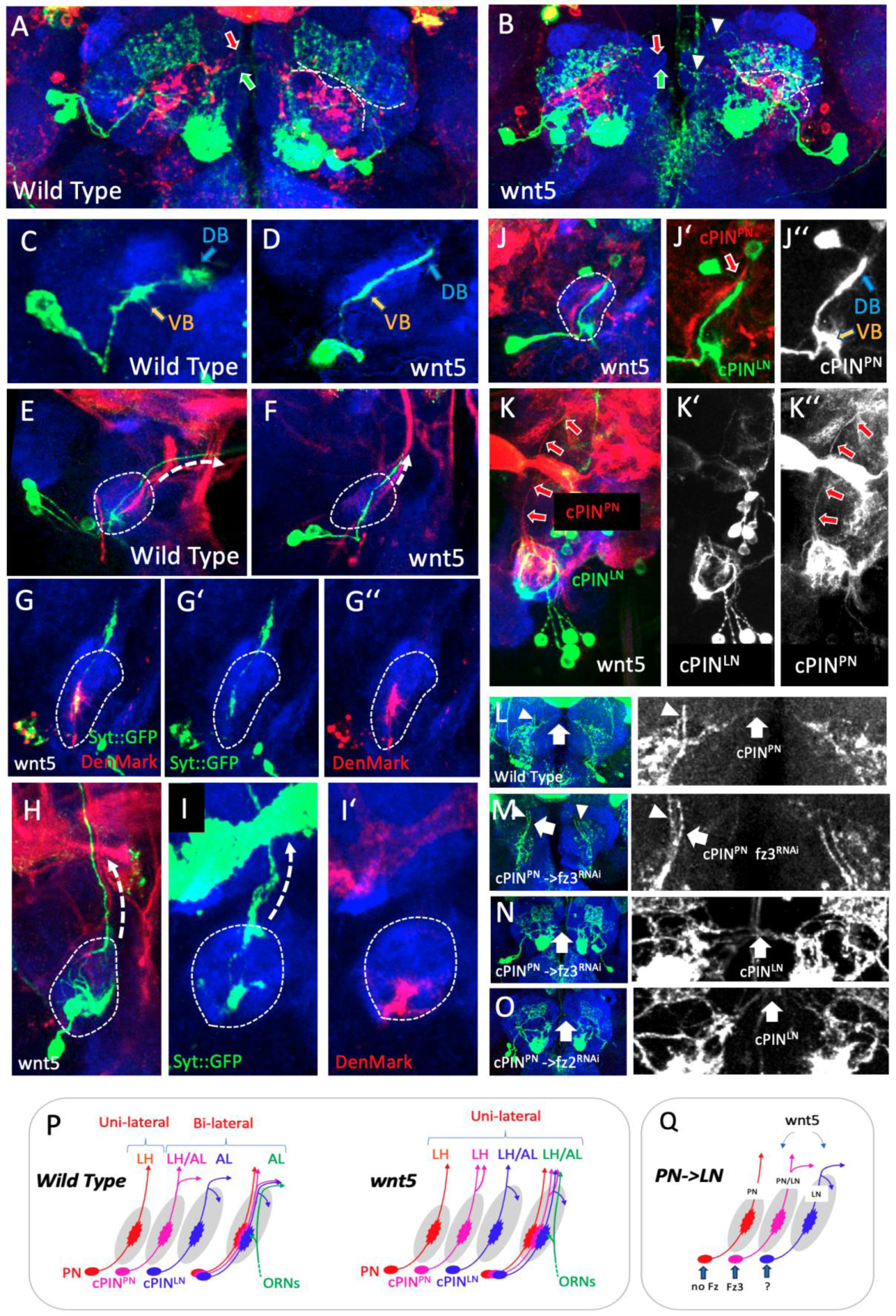
Wnt signaling interferes with a default PN differentiation program to support local cPIN relay. (A-B) The overall dendrite pattern of cPIN^LN5^ and cPIN^PN^ classes is maintained in *Wnt5* mutants. However, loss of *Wnt5* leads to less defined and partially overlapping dendritic fields (dashed lines) and the absence of commissural projections (arrows) with some ectopic dorsally extending fibers (arrowheads). (C-D) In contrast to the ipsilateral branch formation of medial cPINs in wild type, loss of *Wnt5* results in a strong reduction of VB and DB in early pupal development. (E-F) At the stage of contralateral axon extension in wild type, *Wnt5* mutant cPINs project along the PN axon tracts instead of crossing the midline. (G-I) Axon/dendrite identity of cPIN^LN5^ is maintained in *Wnt5* mutants, both during initial outgrowth (G-G’’) and dorsal extension of the axon process (H-I’). (J-K) Medial cPIN^LN5^ axons follow the lateral cPIN^PN^ processes, which also show a reduced branch arborization (J’-J’’). While dorsally extending lateral cPIN^PN^ axons are maintained throughout pupal development (K’), ectopic cPIN^LN5^ axons retract back into the AL (K’’). (L-O) Wnt receptor function in cPIN organization. In contrast to contralateral projection of cPIN^PN^ in wild type (L) the knock-down of *frizzled3* results in a loss of commissural axons, who exits the AL along the main PN tracts (M). Loss of cPIN^PN^ commissural axons does not affect the contralateral extension of cPIN^LN5^ (N, O). (P, Q) Schematics illustrating how the Wnt5 signaling pathway regulates and interferes with the pattern and bilateral projection of cPIN^PNs^ and cPIN^LNs^ neurons during development.

As lateral cPIN^PN^ neurons in wild type also extend along this PN axon tract before projecting a contralateral branch across the midline, we analyzed the interdependency of cPIN classes in *Wnt5* mutants. Colabeling of cPIN^PNs^ and cPIN^LN^ in *Wnt5* mutants showed that in all cases in which cPIN^PNs^ project ectopically along PN axons beyond the normal terminal point, cPIN^LN^ are following this dorsal pathway instead of crossing the midline, resulting in the loss of bilateral cPIN connectivity (Figure 6J). Interestingly, during subsequent steps in *Wnt5* mutant development cPIN^LN^ retract their PN-type axon extension (Figure 6K).

To further test if Wnt5 signaling prevents cPIN axon processes to leave the AL along PNs we targeted the expression of candidate Wnt5 receptors in different cPIN classes. Following the knock-down of Wnt5 candidate receptors (*fz-1/2/3/4*, *drl-1/2* and *vang*) in cPIN^LN^, no significant commissural defects could be detected (Suppl. Figures). In contrast, the reduction of fz3 receptors in cPIN^PN^ pioneer neurons impairs their contralateral projection in 35% of the cases (Figure 6M, n=17). Interestingly, *fz3* knock down in pioneer cPIN^PNs^ does not interfere with cPIN^LN^ contralateral projection, indicating an additional cPIN class which supports cPIN^LN^ connectivity. Together with their polarized axonal-dendritic growth pattern these results indicate a common initial differentiation program shared between unilateral PNs, bilateral PNs and sparse LNs. Different response profiles of polarized neuron processes towards external guidance cues specify their mature olfactory class identity (Figure 6P, Q).

## Discussion

### cPINs as an early integrator of sensory cues

The antennal lobe constitutes the first synaptic integration center for afferent receptor neurons of olfactory, hygro- and thermosensory modality (Marin et al., 2020; Schlegel et al., 2021). Individual glomeruli serve as distinct functional units, each defined by convergence of receptor neurons that share a common receptor identity, for both olfactory and non-olfactory channels (Laissue and Vosshall, 2008; Grabe et al., 2016). Detailed synaptic connectivity between distinct sensory modalities within the antennal lobe has not yet been conclusively analyzed and demonstrated. So far, studies indicate that interconnections are largely restricted to circuits within the same sensory modality, whereas cross-modal integration appears to be mediated predominantly by higher brain centers (Schlegel et al., 2021). Here we show that in addition to their role in supporting fast interhemispheric communication of lateralized olfactory cues, a small class of bilateral interneurons mediate the early ipsilateral convergence of olfactory information with other sensory modalities via lateral AL relay.

Medial cPIN^LN5^ collects olfactory input of ammonia/amine identity and converges with a sensory pathway for humidity channel responsible for the detection of evaporative cooling (Marin et al., 2020). Both environmental humidity levels and a high-protein food sources provide important information for fly behavior, for example in foraging and egg-laying decisions. As many other insects, Drosophila flies are sensitive to humidity changes as their high surface-to-volume ratio makes them vulnerable to desiccation (Kühsel et al., 2017). However, several studies could show that, without an internal physiological state of thirst, Drosophila will prefer an attractive food source over an ideal external humidity (Sayeed and Benzer, 1996; Liu et al., 2007; Jourjine et al., 2016). This is in line with our findings of cPIN^LN5^ inhibiting the positive valence of PN^VP1d^ mediated humidity perception in the presence of ammonia/amine sensation.

Three distinct classes of commissural pioneer interneurons (cPINs) can be distinguished by their input and output characteristics: two primarily mediate intra-modal connectivity, either linking olfactory pathways or integrating temperature and humidity circuits (cPIN^LN31^ and cPIN^PN^), whereas a third class (cPIN^LN5^) uniquely functions as an intermodal bridge, integrating ammonia sensory input and modulating the evaporative cooling pathway. All three cPIN classes provide input onto various PN neuron types in the posterior AL, with dendritic fields not restricted to a single sensory class but collecting information from various other PNs and convey the information to different integration centers, such as the WED, PLP, AVlP and PVL (Marin et al., 2020). These regions seem to integrate various sensory inputs and relay the information to descending motor neurons (Namiki et al., 2018), which is another indicator for fast response pathways.

### cPINs support the establishment of early AL circuitry

Synaptic glomeruli function as distinct processing units within the olfactory system for a linear transfer of olfactory receptor identity (Grabe et al., 2016), in which matching sensory and projection neurons build well-defined input and output subunits. In contrast, different classes of multi-glomerular local interneurons allow more global modulation of olfactory information processing (Olsen and Wilson, 2008; Kazama and Wilson, 2009). Here we characterize the circuit organization and developmental profile of a less-characterized class of olfactory interneurons which combines morphological and functional features of both PNs and LNs. While all cPIN processes are restricted to the AL thereby classifying them as LNs, a highly polarized cellular organization with specific glomerular input and output domains are the main characteristics of glomerular PNs. The developmental profile of cPIN growth and glomerular circuit integration suggests a basic genetic program of PN identity modified at the growth phase to prevent AL exit. The early segregation of pre- and postsynaptic domains of cPIN processes, in which the axonal branch is extending along the main PN fiber tract and the dendritic domain becomes integrated into the ORN class specific protoglomerulus are key features of PN connectivity formation (Figure 3J) (Jefferis et al., 2004). The transition of a PN-like to a LN-like identity is mainly achieved by the rerouting of the axonal branch from the ipsi- to the contralateral AL. Interestingly, these different types of relay neurons follow a defined sequential growth pattern in which first glomerular PNs exit the AL to extend to the ipsilateral LH/MB, followed by a bilateral cPIN^PN^ class with hybrid features of a canonical PN morphology and interhemispheric connectivity and the late projection of bilateral cPIN^LN^ classes. We identified Wnt5 as a critical signal to transform the PN projection identity into a bilateral LN, the cPIN^LN^ class. Before sensory axon arrival, dendrites of cPIN^LN^ together with PN dendrites form a rough spatial map which prefigures the later position of the corresponding AL glomeruli (Figure 4N). While the innervation of the anterior AL precursor neuropil by olfactory sensory neurons creates synaptic glomeruli with strict spatial borders (Gao et al., 2000), other sensory modalities from the antenna seems to lack the capacity of glomerulus induction and PN dendrites maintain their more variable organization (Marin et al, 2020). Early sorting of presynaptic markers in extending cPIN^LN^ indicates an intrinsic mechanism similar to PNs for initial axon/dendrite specification (Figure 5L) (Rolls, 2011). In contrast, the later segregation of input-output domains with cPIN^LN^ dendrites seems to be influenced by ORN axons, as we could observe a directional growth of those dendrites devoid of presynaptic markers towards the peripheral protoglomeruli. This step of glomerulus formation cPIN^LN^ also share with the growth pattern of PN dendrites (Figure 4N).

The Wnt5 signaling pathway has shown to be important for multiple aspects of PN development in the Drosophila olfactory system, especially in the early dendrite patterning (Wu et al., 2014). Interestingly, we do not find significant effects on cPIN^LN^ dendrite spatial organization or intrinsic polarity following the loss of Wnt5. Here, Wnt5 signaling seems to be specific for the redirection of axons to the contralateral AL instead of extending towards the LH/MB, indicating molecular differences in the response properties of cPIN^s^ and PNs. In addition, misrouted cPIN axons in *wnt5* mutants show a subtype specific connectivity in adult: while the cPIN^PN^ class maintains the ectopic dorsal projection to the secondary olfactory processing centers, cPIN^LN^ neurons retract the dorsal axon extension by the end of olfactory system development, indicating different synaptic recognition properties.

### Early sensory processing supports efficient behavioral responses

cPINs were first characterized for their essential role in establishing interhemispheric projections for following olfactory receptor neurons (ORNs), which enables early bilateral integration of olfactory signals in the antennal lobes (Kaur et al., 2019). This bilateral connectivity allows fast flying flies such as Drosophila to rapidly compare inputs from the left and right antennae, producing precise directional responses to transient odor stimuli (Grabe et al., 2016; Mohamed et al., 2019). Such fast integration of sensory cues is particularly critical for small, fast-flying insects with short antennae, where latency in sensory processing can affect proper navigation during foraging (Nawrot et al., 2010).

Besides olfaction, Drosophila is dependent on the rapid integration of not just one, but multiple sensory modalities, including visual, mechanosensory and thermosensory inputs. This enables a precise and rapid navigation in dynamic environments. While higher brain regions such as the mushroom bodies and lateral horn are the main processing center for experience dependent and valence-based integration, it seems that multisensory processing can already occur at early stages of sensory pathways to enable faster sensorimotor integration and transformation (Stein and Stanford, 2008; Wilson, 2013). Such early integration across distinct circuits improves accurate orientation and a stable navigation during flight, supporting the fly to adapt to dynamic environmental conditions. (Currier and Nagel, 2020; Suver et al., 2019).

## Material & Methods

### Drosophila Stocks

Flies were raised on food, cooked in our laboratory food kitchen (10g agar, 70g corn flour, 10g soy flour, 18g dry yeast, 70g malt extract, 35g sugar beet syrup, 4ml propionic acid, 15ml nipagin solution (100g in 1l 70% ethanol)) and kept at 25°C. The fly stocks listed below were used for developmental and adult characterization of the cPINLN5 circuit, including co-labeling of candidate synaptic partner neurons and visualization of pre- and postsynaptic compartments. (BDSC = Bloomington stock center): 10xUAS-mCD8::GFP (BDSC 32184), 13xLexAop::mCherry (3rd chr.: BDSC 52271, 2nd chr.: BDSC 52272), UAS-DenMark, UAS-syt.eGFP (BDSC 33065), UAS-Dscam:GFP (BDSC 66201), cPINLN5 and cPINLN31 (BDSC 49852), cPINPN (cPINLN38; BDSC 48867), DP1m (BDSC 41732), RH50-Gal4 and amt-Gal4 (VM6; stocks kindly provided by Karen Menuz’ lab), VM1 (BDSC 41733), VM4 (BDSC 41735), VP1d (BDSC 41316), VP2 (BDSC 57620), VM6PN (Gal4: BDSC 48506, LexA: BDSC 52446), VP1dPN (BDSC 48795), VP1dPN++ (BDSC 38599), VP1dPN+VL1 (BDSC 47898), VP1dPN+VP4l2 (BDSC 49975), l2PN3t18PN (BDSC 39579).

Neuronal activity was visualized using the UAS-CaLexA reporter line (BDSC 66542), while connectivity patterns were assessed using GFP Reconstitution Across Synaptic Partners (GRASP, BDSC 79040). Behavioral experiments were done with the respective Gal4- and LexA-lines mentioned above to either activate them via UAS-ReaChR (BDSC 53741) and LexAop-ReaChR (BDSC 53747) or inhibit them via UAS-Aurora (BDSC 76237).

To investigate whether known signaling molecules interfere with the correct establishment of the cPIN circuit, we utilized the following lines: wnt5^400^(BDSC 64300), UAS-fz2-RNAi (BDSC 27568), UAS-fz3-RNAi (2^nd^ chromosome: BDSC 66951; 3^rd^ chromosome: BDSC 44468).

### Connectome Analysis

The NeuPrint hemibrain connectome dataset (v1.1; Janelia Research Campus) was utilized to reconstruct and evaluate synaptic connectivity matrices for the neurons of interest and their downstream partners. Morphological skeletons, synaptic locations, and pre- and postsynaptic distributions were visualized and analyzed using the NeuPrint and NeuroNLP software platforms (https://neuprint.janelia.org and https://hemibrain.neuronlp.fruitflybrain.org/).

### Immunohistochemistry

The selected adult flies were anesthetized with CO2 and killed in 96% ethanol. After that, they were washed several times with PBS and then transferred to a drop of PBS (130mM NaCl, 7mM Na2HPO4, 3mM KH2PO4, 2,7mM KCl) on a dissection plate, covered with silicone. Using forceps, the head was opened and the cuticula was removed from the brain. Larvae and pupae were collected and directly dissected. Afterwards they were transferred to Eppendorf tubes filled with a 2% PFA (10ml PBS, 0,8g PFA, 70µl 1M NaOH) solution for a one-hour fixation at 25°C. Then the fixated brains got washed 4 times with 0,3% PBT (PBS + 0,3% Triton X-100) for 15 minutes on the shaker at 25°C. After one hour of washing, the brains got incubated in 10% goat serum solved in 0,3% PBT at 25°C for one hour. Then the appropriate first antibody was applied and incubated over night at 4°C. The next day the first antibody was removed, washed 4 times with 0,3% PBT for 15 minutes on the shaker at 25°C. After that, the appropriate second antibody was applied and again incubated over night for 4°C. On the next day, the second antibody got removed and washed for 4 times with 0,3% PBT for 15 minutes on the shaker at 25°C and mounted on specimen slides.

The mounting was done with fluorescence microscopes to orient the brains properly, in this case mostly anterior to posterior. Afterwards the samples were imaged by a Leica SP5-2 DM6000 confocal microscope. The settings for GFP adjustments were kept the same for the serotonin sensor experiments, so that the endogenous signal can be analyzed and compared afterwards. Finally, the images were processed using the image processing program ImageJ® and IMARIS®.

Antibody staining for different experiments was done as followed: For background staining Anti-Ncad rat (1:10) and anti-rat 647 (1:500) was used to label all neurons in the fly brain. To label the distinct proteins expressed by MCFO, anti-flag mouse (1:2000) and anti-HA rabbit (1:2000), as well as anti-mouse 568 (1:300) and anti-rabbit (1:300) was used for staining.

### Multicolor flipout

To label individual cells within an otherwise broadly expressed Gal4 line, we employed a method developed by Nern et al., (2015). This approach uses four independent constructs, each consisting of a UAS sequence followed by a transcriptional stop cassette and one of four distinct transmembrane proteins that can be detected with specific antibodies. Following stochastic activation, these constructs can be expressed individually or in combination within the same cell, generating unique color combinations and thereby increasing the likelihood of distinguishing single cells from their neighbors. The stop cassette is excised stochastically by a heat-inducible recombinase. Activation of the recombinase is achieved by incubating flies at 37°C, resulting in sparse and random labeling of cells.

### GFP reconstitution across synaptic partners

To visualize synaptic connections, we used targeted-GRASP, a technique based on the reconstitution of split GFP at sites of synaptic contact. In this system, GFP fragment 11 is fused to the presynaptic protein Cacophony, while GFP fragments 1–10 are attached to the postsynaptic protein Telencephalin (Shearin et al., 2018). The membrane-bound nature and compartment-specific localization of these fusion proteins enable the selective labeling of synaptic contacts independently of neuronal activity. Consequently, GFP fluorescence is reconstituted only at sites where the presynaptic and postsynaptic membranes are in close proximity, allowing the visualization of synaptic connections between defined neuronal populations.

### Denmark and synaptotagmin

Under the control of the Gal4/UAS system, distinct neuronal compartments can be visualized using compartment-specific fluorescent markers. Dendrites are labeled in red through the expression of Telencephalin fused to mCherry, whereas presynaptic terminals are labeled in green by expressing Synaptotagmin fused to GFP (Nicolai et al., 2010). This approach enables the simultaneous visualization of dendritic and presynaptic regions within neurons of the selected Gal4 line.

### Optogenetics

For the optogenetic behavioral experiments, age-matched adult flies (3–4 days old, both sexes) were reared in complete darkness at 25°C on food supplemented with all-trans-retinal to enhance light sensitivity. To reduce visual confounds that could influence behavior, all handling was performed in darkness under dim red light, which is minimally perceived by the flies.

Flies were transferred from incubator vials into empty vials connected to the optogenetic apparatus and moved into the elevator compartment. They were allowed to acclimate for 60 seconds just above the decision point. Subsequently, green LED illumination was activated, and the elevator was lowered to initiate the trial. The flies were then given 20 seconds to explore the arena and make a choice. After this period, the elevator was raised, separating the animals into either the test arm (light), the control arm (dark), or the central elevator region.

A Preference Index (PI) was calculated by subtracting the percentage of flies in the test arm from the percentage in the control arm. Flies remaining in the central elevator were classified as neutral; their numbers were evenly distributed between the test and control groups before calculating final percentages.

For neuronal manipulation, we used an optogenetic strategy based on Inagaki et al. (2014), employing UAS-ReaChR and LexAop-ReaChR in combination with Gal4- and LexA- driver lines to activate target neurons with green light (530 nm). In addition, UAS-Aurora (Wietek et al.) was used to silence targeted neurons via red-light illumination (585 nm).

### Statistical analysis

Statistical analyses and graphical illustrations were performed using GraphPad Prism 9 software. Statistical significance across genetic groups was evaluated using a One-Way ANOVA followed by Tukey’s post-hoc multiple comparisons test (or Kruskal-Wallis test where appropriate). Error bars indicate the mean ± SEM. Significance thresholds are indicated as follows: p < 0.05 (*), p < 0.01 (**), p < 0.001 (***). Each dot in the figures represents an independent experimental replicate.

